# A Spiking Neural Network Model of Rodent Head Direction calibrated with Landmark Free Learning

**DOI:** 10.1101/2022.02.01.478640

**Authors:** Rachael Stentiford, Thomas C. Knowles, Martin J. Pearson

## Abstract

Maintaining a stable estimate of head direction requires both self motion (ideothetic) information and environmental (allothetic) anchoring. In unfamiliar or dark environments ideothetic drive can maintain a rough estimate of heading but is subject to inaccuracy, visual information is required to stabilise the head direction estimate. When learning to associate visual scenes with head angle, animals do not have access to the ‘ground truth’ of their head direction, and must use egocentrically derived imprecise head direction estimates.

We use both discriminative and generative methods of visual processing to learn these associations without extracting explicit landmarks from a natural visual scene, finding all are sufficiently capable at providing corrective signal. Further, we present a spiking continuous attractor model of head direction (SNN), which when driven by ideothetic input is subject to drift. We show that head direction predictions made by the chosen model-free visual learning algorithms can correct for drift, even when trained on a small training set of estimated head angles self-generated by the SNN. We validate this model against experimental work by reproducing cue rotation experiments which demonstrate visual control of the head direction signal.

## Introduction

As we move through the world we see, touch, smell, taste and hear the environment around us. We also experience a sense of our own self-motion through our vestibular system, which enables us to keep balance and to maintain an internal estimate of our location and heading, or pose in the world. Any drift in pose estimate incurred through the integration of self-motion cues alone (as we walk with our eyes closed for example) is quickly corrected when we open our eyes and recognise familiar features in the environment. This approach to self localisation has been adopted in many fields of engineering that require an accurate and persistent pose estimate to operate effectively such as mobile robots and augmented reality devices.

The relative contributions of self-motion (ideothetic) and external sensory cues (allothetic) to the firing properties of ’spatial cells’ in rodents has been extensively investigated in neuroscience (see below for review). The integration of ideothetic cues provides the animal with a rapid and constant estimate of pose. This estimate not only aids navigation in the absence of allothetic cues, but is also a learning scaffold to associate pose with novel visual scenes. This second function has received little attention in prior models which often use the ground truth pose of a learning mobile agent to associate sensory view rather than the drift prone estimate provided from ideothetic cues. In mobile robotics this problem is addressed in the research field known as Simultaneous Localisation And Mapping (SLAM) with myriad solutions proposed, each with their own advantages and limitations based on environmental, sensory and computational constraints. In this study we are interested in modelling how this problem has been solved in the mammalian brain.

The problem of accumulative error in ideothetic cue integration implies that head direction estimates from early explorations of a novel environment should be more reliable than later explorations. Therefore, earlier experiences of an environment should be used to learn associations between heading and visual scenes, to correct for drift as the animal later explores the same environment. From a learning perspective this puts constraints on the size and richness of training sets. This bi-directional learning problem is investigated here through a series of controlled experiments using a simulated mobile robot within a virtual environment. Our contribution is to model the head direction cell system using populations of spiking neurons, translating angular head velocity cues into a spike encoded representation of head direction. To anchor head directions to allothetic cues, we have trained three different model-free visual learning algorithms: Convolutional Neural Network (CNN), Variational Auto-Encoder (VAE) and Predictive Coding Network (PCN), to associate distal natural visual scenes with the spike based representation of head direction. We demonstrate that all three models are capable of correcting the drift in pose estimate from purely ideothetic cue integration even when trained on small self-generated training sets. This is evaluated further through cue conflict experiments to reveal similar characteristics of the model performance as recorded in rodents.

### Rodent head direction cell system

Neural correlates of position ([O’Keefe, 1976, Hafting et al., 2005]), environmental boundaries ([Lever et al., 2009]), heading ([Taube et al., 1990]), speed ([Kropff et al., 2015]) and numerous other spatial measures (see [Grieves and Jeffery, 2017] for review) have been extensively studied in the rodent brain and remain an active topic in neuroscience research. Of these, Head Direction (HD) cells which exhibit high firing rates only in small arcs of head angle, appear simplest and have been a popular target for modelling.

Most models of head direction use a continuous attractor where a sustained bump of activity centred on the current heading is formed and maintained through interactions between excitatory and inhibitory cells. Many rely on recurrent excitatory collaterals between cells in the Lateral Mammillary Nuclei (LMN) ([Zhang, 1996, Page and Jeffery, 2018]), however anatomical data show no evidence of this type of connection ([Boucheny et al., 2005]). Although head direction cells have been found in many brain regions, including Anterior Thalamic Nuclei (ATN; [Taube, 1995]), Retrosplenial cortex ([Cho and Sharp, 2001]), Lateral Mammillary nuclei (LMN; [Stackman and Taube, 1998]) and Dorsal Tegmental Nucleus (DTN; [Sharp et al., 2001]; see [Yoder et al., 2011] for review), generation of the head direction signal is thought to be in the reciprocal connection between LMN and DTN ([Blair et al., 1999, Bassett and Taube, 2001]). As the DTN sends mainly inhibitory connections to the LMN, attractor networks exploiting connections between two populations of cells appear more biologically plausible ([Boucheny et al., 2005, Song and Wang, 2005]).

### Control of HD by self-motion cues

Self-motion cues can be derived directly from the vestibular system but also from optic flow (Arleo et al 2013) and motor efference copy. Disrupting vestibular input to head direction cells abolishes spatial firing characteristics and impacts behaviours which rely on heading ([Yoder and Taube, 2009, 2014]). Cells sensitive to Angular Head Velocity (AHV) have been recorded in several regions including the DTN ([Sharp et al., 2001, Bassett and Taube, 2001]). These cells are either sensitive to AHV in a single direction (clockwise or anticlockwise; asymmetric AHV cells) or the magnitude of AHV regardless of direction (symmetric AHV cells). Methods of moving the bump of activity on the ring attractor to follow head movement rely mainly on asymmetric AHV input. Bump movement is achieved either through imbalance between two populations of cells in the attractor network ([Bicanski and Burgess, 2016, Boucheny et al., 2005]), or via conjunctive cells which fire strongly as a function of both AHV and head direction ([Sharp et al., 2001, McNaughton et al., 2006]). However using imprecise self motion cues in the absence of vision results in drift in the preferred firing direction of head direction cells ([Stackman et al., 2003])

### Visual control of HD

Although HD cells still show some directional sensitivity in the absence of visual cues or novel environments ([Goodridge and Taube, 1995, Taube and Burton, 1995, Goodridge et al., 1998, Stackman et al., 2003]), vision is clearly an important factor for stabilising the head direction system. During development, head direction cells have much sharper tuning curves after eye opening (Tan et al 2015), but may use other types of allothetic information such as tactile exploration of corners of the environment with whiskers to stabilise head direction before eye opening ([Bassett et al., 2018]). Even in unfamiliar environments, visual information helps to stabilize head direction, suggesting ongoing learning of visual landmarks ([Yoder et al., 2011]). In familiar environments, the preferred firing directions of head direction cells become entrained to visual features and will follow these cues over self-motion signals ([Taube and Burton, 1995]). When environmental cues are rotated, preferred firing directions of many cells also rotate through the same angle, resulting in ‘bilateral’ preferred firing directions, Page and Jeffery (2018) suggest these bilateral cells may be useful for assessing the stability of environmental landmarks. Some head direction cells won’t follow these big conflicts in cue location, suggesting multiple populations of head direction cells that are more or less strongly controlled by allothetic input ([Dudchenko et al., 2019]). This visual control of head direction begins at the LMN ([Yoder et al., 2015]), stabilising the head direction signal at its origin. Both the postsubiculum (PoS) and retrosplenial cortex (RSC) are likely candidates for delivering this visual information to the LMN ([Taube et al., 1990, Vann et al., 2009]). There are two main methods of using visual information to control the head direction bump position. The first is to use visual information to calibrate the model of AHV, whether that be by detecting error between the estimated head angle and the expected head angle based on the visual cue ([Kreiser et al., 2020]), or using a combination of strategic behaviour and landmark tracking to match the AHV model to the movement of the cue within the visual field ([Stratton et al., 2011]). The second method is to influence the head direction bump position directly by exploiting the attractor dynamics and injecting current into the new bump position. This could simply use Gaussian inputs into the ring attractor at determined positions ([Song and Wang, 2005]), or by representing features in multiple ‘landmark bearing cells’, learning the association between head angle and visible features, and feeding back expected head direction onto the head direction cells ([Yan et al., 2021]). A combination of these two methods of visual control is probably the answer to accounting for environmental or body changes in the real world, in this study we begin with directly influencing the bump position.

### Models of visual input

In models of head direction stabilised by visual input, the visual data used is often very controlled. For example, adopting ‘visual cells’ that fire at specific head angles without any real visual data ([Song and Wang, 2005]) and assume visual processing is performed somewhere upstream. Where true vision is used (captured by cameras), one or more cues positioned in the environment such as coloured panels ([Yan et al., 2021]) or LEDs ([Kreiser et al., 2020]) are identified, mapped to a ‘visual cell’ and learning mechanisms associate this cue with a head angle ([Bicanski and Burgess, 2016]). Natural visual scenes are much more complex and information rich than bold homogenous cues, with this richness making real-world landmark identification more difficult. The question remains, how does a visual scene become useful for maintaining head angle? In this work we use natural scenes, projected onto a sphere around a simulated robot (see figure 1). We assume this visual information is distal and invariant to translation. We show that both generative and discriminative model-free learning algorithms can be used to predict head angle from natural visual information and correct for drift in a spiking continuous attractor model of head direction cells, without the need to identify specific landmarks in the environment.

**Figure 1:**
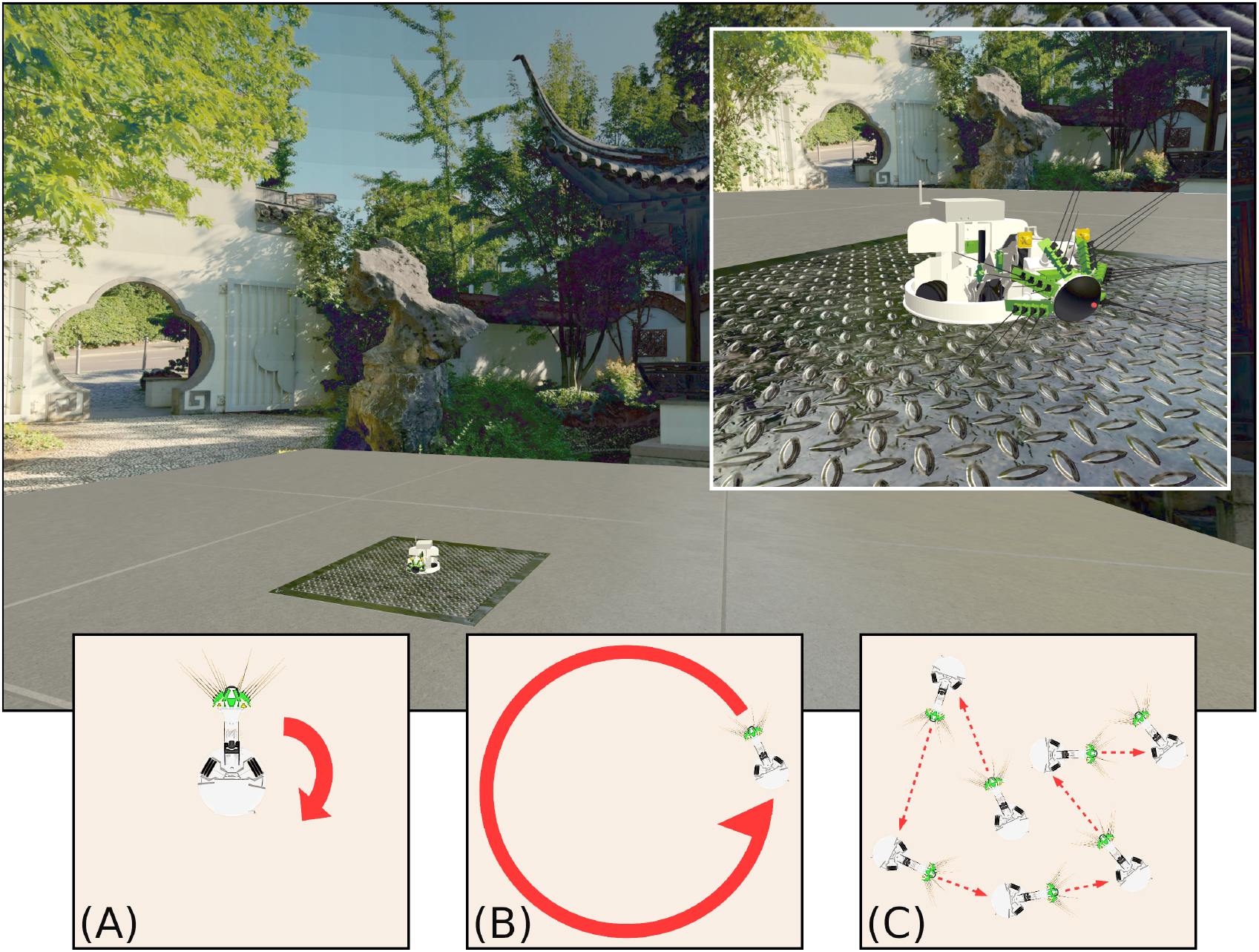
The WhiskEye robot used to capture visual and odometry data sets as it moved within the simulated environment of the NeuroRobotics Platform. The Natural scenery (a panorama of the Chinese Garden in Stuttgart) was projected onto the inside surface of a sphere that surrounds the platform on which WhiskEye can move. The behaviours expressed by WhiskEye during capture of the data sets analysed in this study are referred to as **(A)** rotating, **(B)** circling, and **(C)** random walk

#### On the Discriminative-Generative dichotomy

Machine Learning approaches broadly draw from two paradigms. The first is a discriminative paradigm; models which aim to partition the incoming data samples along meaningful boundaries, often building a hierarchy of increasingly abstract representations or increasingly broad sub-spaces, extracting meaning through a bottom-up feedforward process. An example would be Decision Trees, which learn to partition data through recursive splitting on the data space by simple if-else rules ([Breiman et al., 1983]).

The second is a generative paradigm, which approaches the problem in the opposite direction. A generative model, often also a probabilistic model, aims to instead learn the capability of generating appropriate data samples like the training data in the appropriate contexts. Like a discriminative model, it tries to uncover abstract features in the data, but instead incorporates this into a model of latent features, refining its hypotheses about the underlying causes of the sensory data it is receiving. An example would be Gaussian Mixture Models, which model the problem space as a family of Gaussians with different parameter values [Reynolds, 2009].

Although details between each model vary considerably, the broad trend is that discriminative models are faster than their generative counterparts, but can only work within the bounds of the data they are provided. With the data space’s dimensionality being potentially unlimited, this still provides a huge amount of capability, but a training set that does not adequately reflect the data space can lead to nonsensical outputs. Generative models, on the other hand, typically tend to be slower to categorise and slower to learn. However, by generating samples from a model of latent causes of the data, they are not limited by their inputs and can produce very different predictions from the data they are provided. For the case of well-defined and well-bounded problems, this is often surplus to requirements, but for many situations, such as with unfamiliar or incomplete data, this can be beneficial.

Many algorithms make use of elements of both. For example, the Variational Autoencoder ([Kingma and Welling, 2013]) has hidden layers that extract features from the data in a discriminative way, and use these features to train a multidimensional Gaussian space, the output of which is decoded by another discriminative layer stack to produce a sensible reconstruction of the input. A Generative Adversarial Network ([Goodfellow et al., 2014]) takes this even further, using a generative and discriminative network in a collaborative competition to produce ever-better data samples. In neuroscience, particularly regarding the visual system, aspects of cortical function have been explained as both a discriminative and generative model, with exactly where and how these approaches synthesise together an active area of research ([DiCarlo et al., 2021]); neural codes originally found in hippocampal work have been hypothesised as a unifying computational principle ([Yu et al., 2021]; see also [Hawkins et al., 2019]).

In this study we remain agnostic to the debate, instead choosing to evaluate a mix of generative and discriminative algorithms for generating predictive head direction signals from allothetic (visual) cues. As a purely generative model, a Predictive Coding Network based on MultiPredNet [Pearson et al., 2021], originally from [Dora et al., 2018]; as a hybrid model, a modification of the JMVAE from [Suzuki et al., 2017]; and as a purely discriminative model, a Convolutional Neural Network ([Bengio and Lecun, 1997]).

## Methods

### Experimental apparatus

WhiskEye is a rat-inspired omnidrive robot with RGB cameras in place of eyes and an array of active whisker-like tactile sensors as shown in Figure 1. In this study only the visual frames from the left camera were considered. A simulated model of WhiskEye was integrated into the Human Brain Project’s NeuroRobotics Platform (NRP) as part of prior work ([Pearson et al., 2021], [Knowles et al., 2021]). The NRP integrates robot control and simulation tools, such as ROS and Gazebo, with neural simulators, such as NEST ([Falotico et al., 2017]). By running these in a synchronised way on a single platform, simulated robots can interact live with simulated neuron models and allows for experiments with biomimetic and bio-inspired systems. Behaviours can also be specified in more controlled ways using the familiar ROS framework whilst capturing data from both the robot and neural simulators for off-line analysis.

Using the NRP allows arbitrary visual scenes to be constructed within the environment. The visual scene in this experiment consisted of a concrete-textured floor for WhiskEye to move on, surrounded by an invisible collision mesh to contain the robot in the environment, and with an outer sphere to display the background. The sphere was made large enough so that translation had no perceptible effect on the visual scene; therefore, barring the concrete floor, all visual cues could be considered distal. Within this environment, WhiskEye executed three different behaviours; rotating on the spot, circling around the centre of the environment and a random walk (illustrated in panels A-C of Figure 1). This provided the odometry and visual data with which to validate the performance of each model.

### Spiking neural network model of Head direction cell system

The Head Direction system model is a spiking neural network (SNN) model written in pyNEST (2.18; Eppler). All cells are simulated using the standard leaky integrate-and-fire neuron model (iaf psc alpha) which uses alpha-function shaped synaptic currents. The simulation timestep was set to 0.1 ms for high accuracy with synaptic delay of 0.1 ms. The network is composed of four equally sized rings of neurons: 180 Lateral Mammillary Nuclei (LMN) cells, 180 dorsal tegmental nuclei (DTN) cells, 180 Clockwise conjunctive cells and 180 Anticlockwise conjunctive cells. Constant input current of 450 pA to all LMN neurons results in spontaneous firing at a rate of 50 spikes per second prior to inhibitory input from the DTN.

Attractor dynamics emerge through reciprocal connections between cells in the excitatory LMN population and inhibitory DTN population. Each LMN cell connects to a subset of DTN neurons with declining synaptic strength as a function of distance (Figure 2A). Reciprocal inhibitory connections from the DTN to LMN cells are arranged with synaptic strength decreasing as a function of distance offset by a constant (mu). This arrangement provides inhibitory input to the cells surrounding the most active LMN cell, producing a single stable bump of activity.

**Figure 2:**
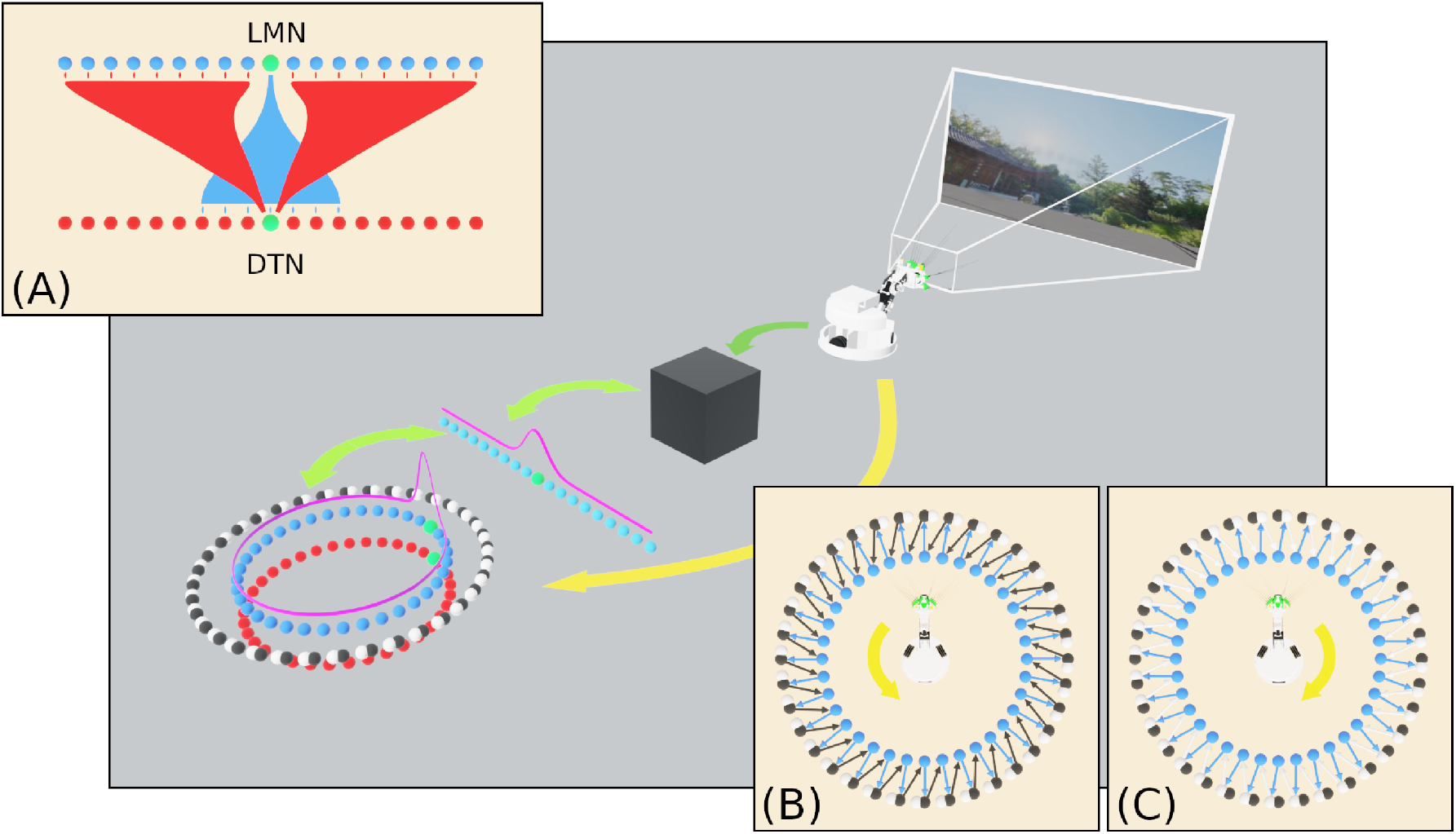
An overview of the SNN model of head direction, the information flows and experimental setup of this study. Visual data from WhiskEye (dark green arrow) and the estimated head angle from the SNN (light green arrows) are used to train one of several model-free learning algorithms (black box). Once trained, these algorithms return head direction predictions (represented as the activity in the array of light blue neurons) that are mapped one-to-one with HD cells in the LMN ring (dark blue neurons) to correct for drift in the spiking HD ring attractor model. **(A)** Excitatory and inhibitory projections between the LMN and DTN respectively for the current most active cell (green neuron). Attractor dynamics emerge from this connectivity to maintain the bump in a stable position in the absence of idiothetic input. **(B)** Connectivity between anticlockwise conjunctive cells (black neurons) and head directions cells offset by one cell anticlockwise. With coincident head direction and angular velocity input (yellow arrow) these cells drive the bump clockwise around the ring. **(C)** Connectivity between clockwise conjunctive cells (white neurons) and head directions cells.

LMN and DTN cells are arranged as rings for the purpose of defining synaptic strength based on distance, this gives the attractor network periodic boundaries. The synaptic weight between each pair of LMN and DTN cells is defined as follows:

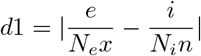

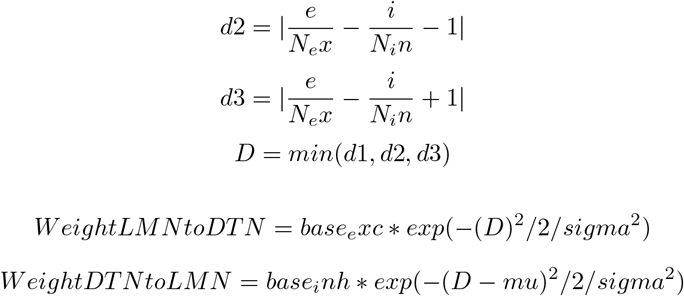

In the absence of input from the two conjunctive cell populations, this bump of activity remains stationary. The initial position of the activity bump is produced by applying a 300 pA step current for 100 ms to a nominal LMN cell.

In order to track head direction based on the Angular Head Velocity (AHV) the two populations of conjunctive cells are connected one to one with a LMN cell, shifted one cell clockwise or anticlockwise from the equivalently positioned neuron (Figure 2B,C). Angular velocity of the head was determined by taking the first derivative of the head position captured from the simulated WhiskEye at a rate of 50 Hz. Positive velocities indicated anticlockwise head movements and negative velocities indicated clockwise head movements. AHV was converted to current (Ivel) using the following formula:

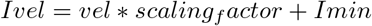

Where Imin = 150 pA, scaling factor = 3500, vel = angular head velocity. A step current generator supplies this current to the respective conjunctive cell population. LMN cells also connect one to one with the equivalent conjunctive cell in both the clockwise and anticlockwise populations. Spiking activity occurs in the conjunctive cells with coincident AHV and LMN spiking input. Conjunctive cell input causes movement of the attractor network activity bump to follow head movement.

#### Allothetic correction

Head direction predictions from the visual learning models (see below) are mapped one to one onto the respective head direction cells as current injections. Negative values in predictions are removed by adding the smallest value in the dataset, then prediction values are scaled by a factor of 10 and supplied to the HD network as a direct current injection. This simple method allows predictions which are smaller in magnitude to have less impact on the bump location. However imprecise predictions, that may have multiple peaks or a broader shape will lead to current input into more cells compared to a perfectly reconstructed Laplacian (see discussion).

### Analysis of network output

To compare the spiking network bump position to the ground truth, the most active cell in each 40ms bin is found. The difference between the estimated head angle and the ground truth was used to show how accumulation of drift over time, with total error measured as Root Mean Squared Error (RMSE). To illustrate drift in the estimated head angle, the preferred firing direction was plotted using firing rate as a function of head angle in polar tuning curves. The ground truth head direction at each spike time was collected into bins (6 degrees), for the first and the third minute, to show changes in preferred firing direction over time. Statistical tests were performed using SPSS statistics software. Differences between ideothetic only and three correction methods were compared using a one-way ANOVA combined with the Tukey HSD post-hoc test. Random predictions, a list of random numbers between zero and the maximum value from the true predictions, generated using a uniform distribution, were used to show that reductions in drift were not due to arbitrary current input.

### Artificial cue rotation

To investigate the control of allothetic cues over the head direction cell signal, we reproduced cue rotation experiments used in rodents ([Taube and Burton, 1995]). To supply current as if environmental cues were rotated by *π*/2 radians, the predictions were manipulated by taking either the first 45 or 135 prediction values and shifting them to the end of the 180 element prediction, producing an artificial rotation. This rotation was applied for 30 seconds after 1 minute of standard predictions.

### Model-free learning algorithms applied to allothetic cue recall

#### Predictive Coding Network (PCN)

This network was a modified version of the MultiPredNet [Pearson et al., 2021] which was developed and evaluated for visual tactile place recognition. Here the 3 modules that made up the original network (visual, tactile and multisensory modules) were re-purposed as visual, head-direction and multisensory modules. Compared to the conventional feedforward architectures of other algorithms, the PCN relies on feedback connections towards the input data. For each sample, the PCN outputs a prediction from its latent layer that passes through the nodes of the hidden layers to the input later. The weights between each pair of layers transform the prediction from the upper layer into a prediction of the lower layer’s activity. At each layer, the prediction from the layer above is compared to the activity at the current layer and the difference (error) calculated. Weights between layers are then updated locally according to their prediction errors. This eliminates the need for end-to-end backpropagation and increases bioplausiblity. Several network topologies were trialed; the best performing network had direct odometry input into the multimodal latent layer, hence the lack of hidden layers in that stream (see table 1 for summary).

**Table 1:** Model parameters for PCN, VAE and CNN. PCN training epochs are low due to number of inner ‘cause epochs’ (50 for training, 500 for inference) that increase training time, though are strictly a modification of the learning rule rather than extra training epochs. Note that the visual layers for the CNN are 2D Convolution Layers with the kernel shape in brackets. Training epochs vary but all networks were trained to convergence. Hebbian learning is as per [Dora et al., 2018], Backpropagation as per [Chollet et al., 2015]

| Parameter | Values |  |  |
| --- | --- | --- | --- |
|  | PCN | VAE | CNN |
| Visual Input Size | 10800 | 10800 | 10800 |
| Visual Hidden Layers Shape | [1000, 300] | [1000, 300] | [64(4,4), 32(2,2)] |
| Odometry Input/Output Size | 180 | 180 | 180 |
| MSI Layer Shape | 100 | [50, 50] | N/A |
| Training Epochs | 200 | 5000 | 50 |
| Learning Rule | Hebbian | Backprop | Backprop |
| Optimiser | N/A | SGD | Adam |

#### Multi-modal Variational Autoencoder (VAE)

Based on the [Suzuki et al., 2017] Joint Multimodal Variational Autoencoder, the VAE works by compressing inputs via hidden layers of decreasing size, encoding inputs into a bifurcated joint multimodal latent space representing the means and variances of Gaussians. These means and variances are used to generate normally distributed random variables, which are then passed through an expanding set of hidden layers to decode the latent Gaussian output into the same shape and structure as the input data. This encoder-decoder system is trained via conventional error backpropagation, comparing the decoded output to the ‘ground truth’ input and adjusting weights accordingly, with the addition of a KL-Divgergence term to penalise divergence from a *μ* = 0 Gaussian. As with PCN, the best performing network had no hidden layers between odometry input and the latent layers, so these were removed from both the encoder and decoder halves of the network.

#### Convolutional Neural Network (CNN)

As a discriminative network, the CNN has only a visual stream to process, with an output layer the same shape as the odometry data. Unlike the other two networks, the CNN has no latent space to condition and operates purely as a encoder, transforming visual scenes to their appropriate head direction output, with weights updated using conventional backpropagation. It is also the only network that is designed specifically for processing images, with strong spatial priors implicit in the way it processes visual scenes, analysing small areas of the image in parallel via convolutions to produce translation-invariant image features. As the problem exists within a small, bounded space in both the visual and odometry domains for this experiment, the larger benchmark CNNs — AlexNet ([Keshavarzi et al., 2021]), ResNets ([He et al., 2016]) etc. — were not required. Instead, a lightweight, purpose-built CNN was created and optimised using Keras Auto-tuner ([Chollet et al., 2015]).

## Results

### Head direction cell like firing properties

Cells in the LMN, DTN and conjunctive cell populations all showed directional firing specificity as observed in the the rodent brain. Figure 3A shows firing rate as a function of head direction from the equivalent cell in each of the LMN, DTN and conjunctive cell rings. The preferred firing direction of these cells is taken at the peak firing rate and the total range of angles each cell fires over is the directional firing range. The average directional firing range of cells in the LMN was 1.035 +/− 0.011 radians, DTN 4.812 +/− 0.012 radians and conjunctive cells 1.034 +/− 0.011 radians (Figure 3B). This is consistent with the directional firing range of DTN head direction cells in rodents (109.43 +/− 6.84; Sharp et al 2001) being greater than the directional firing range of LMN head direction cells (83.4 degrees; [Taube et al., 1990]).Figure 3C shows firing rate as a function of angular head velocity for one example conjunctive cell, that has similar form to asymmetric AHV cells recorded in the DTN ([Bassett and Taube, 2001])

**Figure 3:**
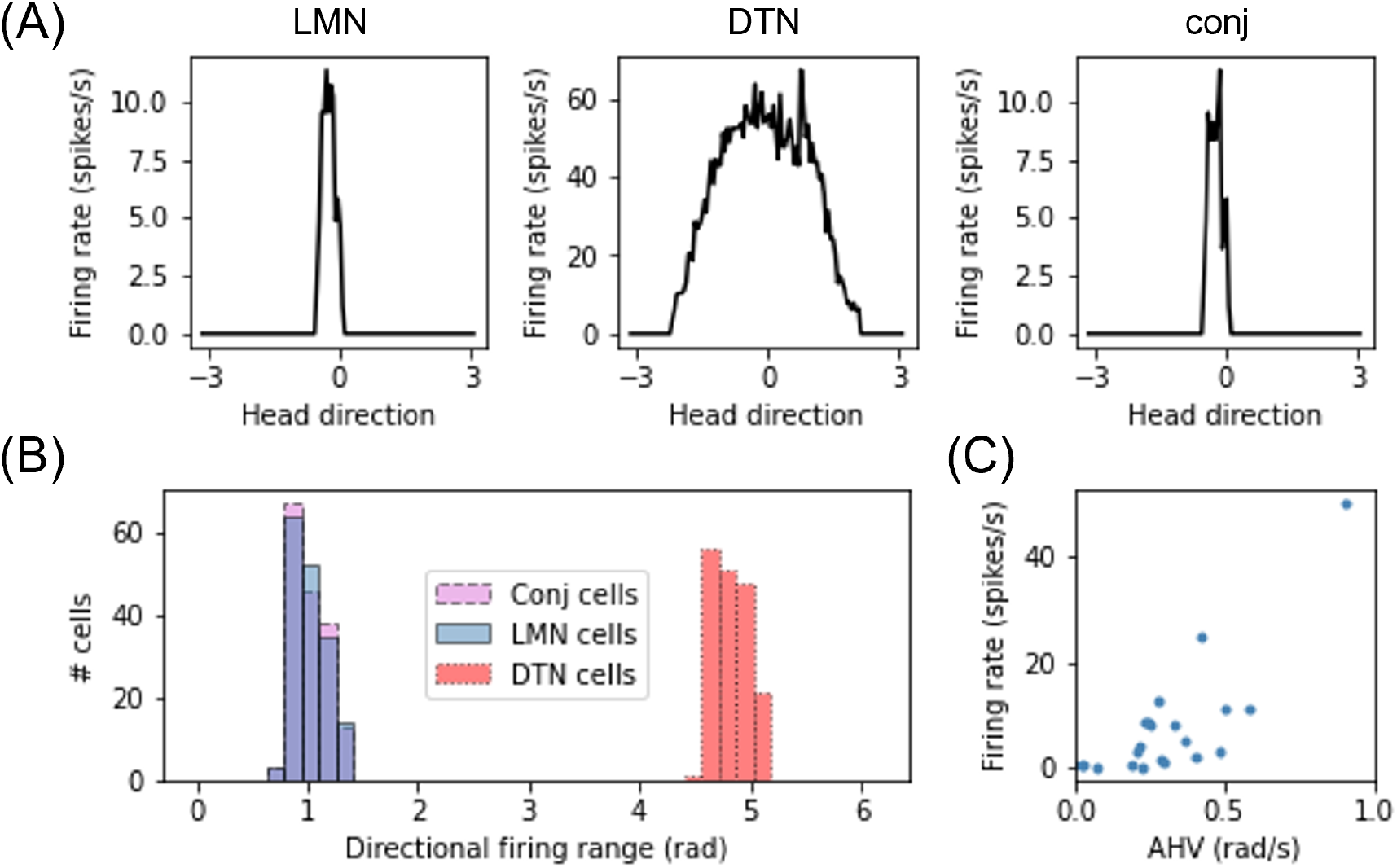
Cells from the DTN, LMN and conjunctive cell populations show HD like firing characteristics. **(A)** Preferred head angle of one DTN, LMN and conjunctive cell expressed as firing rate as a function of head angle, showing strong directional selectivity in the LMN and conjunctive cell and broader directional selectivity in the DTN cell. **(B)** Histogram of the directional firing range of cells in each population, showing broader directional firing in DTN cells. **(C)** Firing rate as a function of angular head velocity (AHV) from one example conjunctive cell.

### Preferred firing direction of cell drift with only ideothetic drive

Ring attractor dynamics which emerge from reciprocal connections between LMN and DTN cells maintain a stable bump of activity centered on the current estimate of head direction. When movement of the bump is driven only by idiothetic angular velocity input from the two conjunctive cell rings, the preferred direction of head direction cells drifted over time. Figure 4A shows the ground truth (grey) and estimated head direction (blue) over time with the WhiskEye robot rotating on the spot. The difference between the ground truth and estimate grows over time (Figure 4B), ending with a maximum difference of 1.65 radians after 3 minutes (RMSE = 1.02 radians). Firing rate as a function of time for 3 LMN cells in the first minute vs the 3rd minute are shown in Figure 4C. The shift in preferred direction of these head direction cells from the first minute to the third minute was 0.896 +/−0.181 radians. However, the RMSE over the first full revolution was fairly low (0.09 radians).

**Figure 4:**
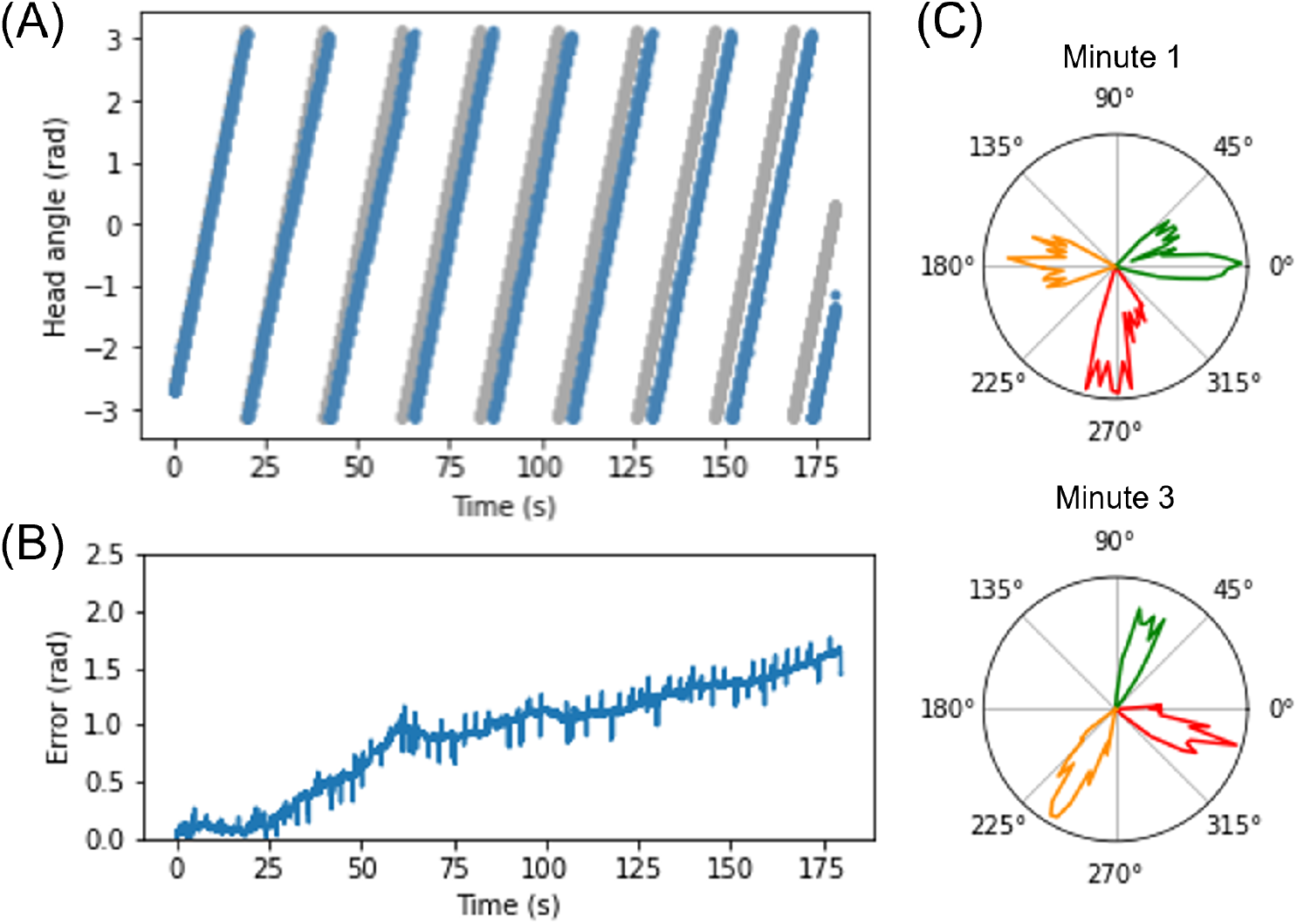
Plots showing drift in the head direction estimate over time. **(A)** Ground truth head angle (grey) and the estimated head angle (blue) from the SNN as the WhiskEye rotates on the spot. Over time the estimate gets further from the ground truth. **(B)** Error measured as the magnitude of the difference between the estimated angle and ground truth increases over time. **(C)** Preferred firing directions of three cells (red, orange and green) in the first vs third minute of the simulation, showing a change in preferred firing direction for all three cell of approximately 70 degrees.

### Predicting head direction using model-free learning algorithms

Rodents use allothetic information such as vision, to counter this drift in head estimate. This requires forming associations between visual scenes and the current head angle, so that the estimated head angle can be corrected when this visual scene is experienced again. Head direction predictions made by three models trained on head direction/vision pairs are not equally structured. As seen in Figure 5, the discriminative CNN is far superior at generating a smooth Laplacian reconstruction, closely approximating the ground truth equivalent. The VAE reconstructions consisted of many competing peaks of varying heights, whilst the PCN shows qualities of both, maintaining a Laplacian-esque area of the distribution with noise increasing as distance from ground truth head direction.

**Figure 5:**
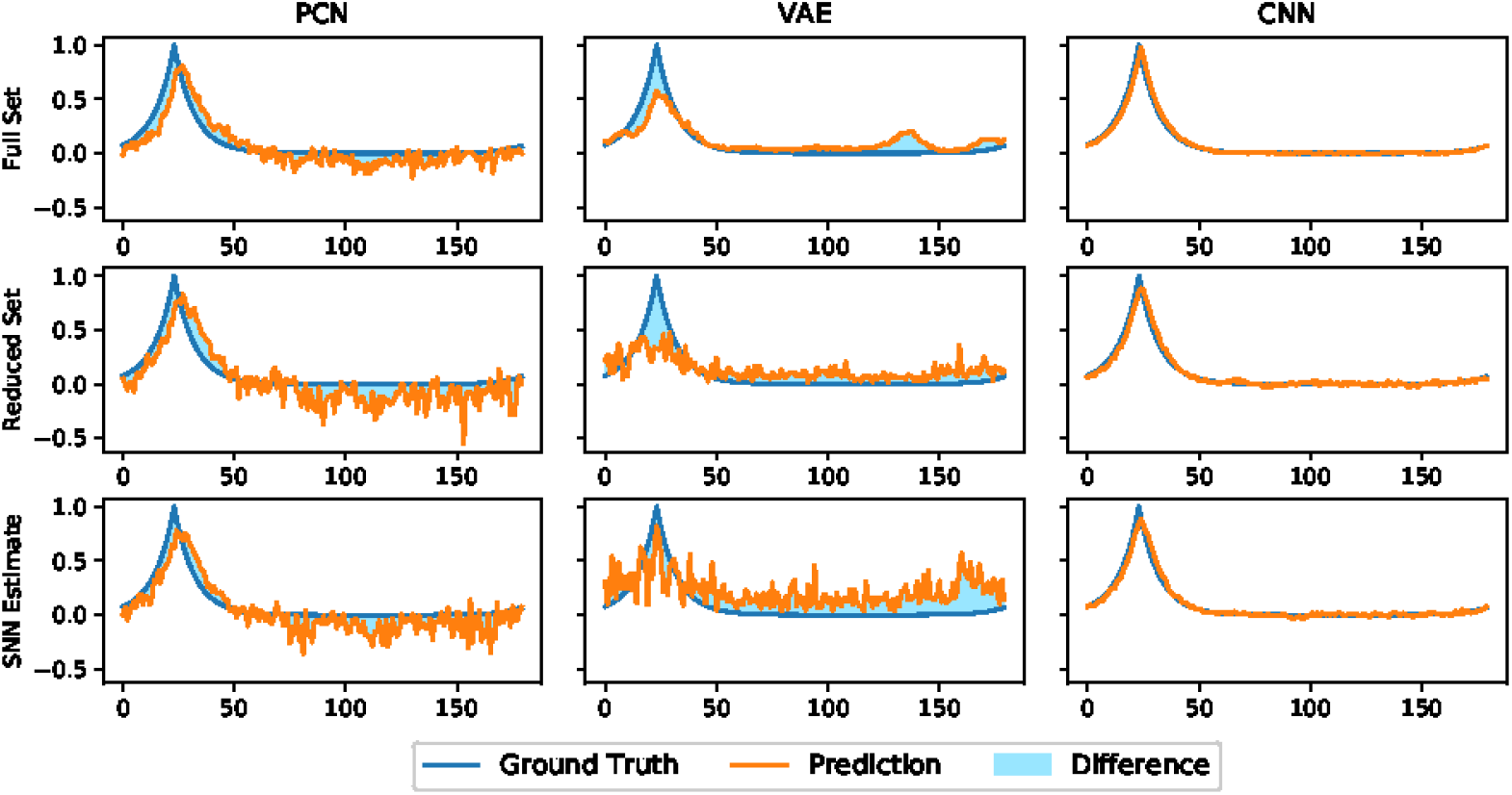
Representative reconstructions of head direction predictions inferred by each algorithm (orange) at a given ground truth head direction (Blue). The shaded difference between the two curves illustrates magnitude of the Root Mean Squared Error, which is inversely proportional to the quality of the reconstruction.

Figure 6 shows the reconstruction error (mean RMSE) for binned views of the visual scene taken from WhiskEye during the rotating behaviour. Although there are variations in the error, the performance remains nominally uniform for all head angles, suggesting that the models are not favouring particular features for head direction estimate.

**Figure 6:**
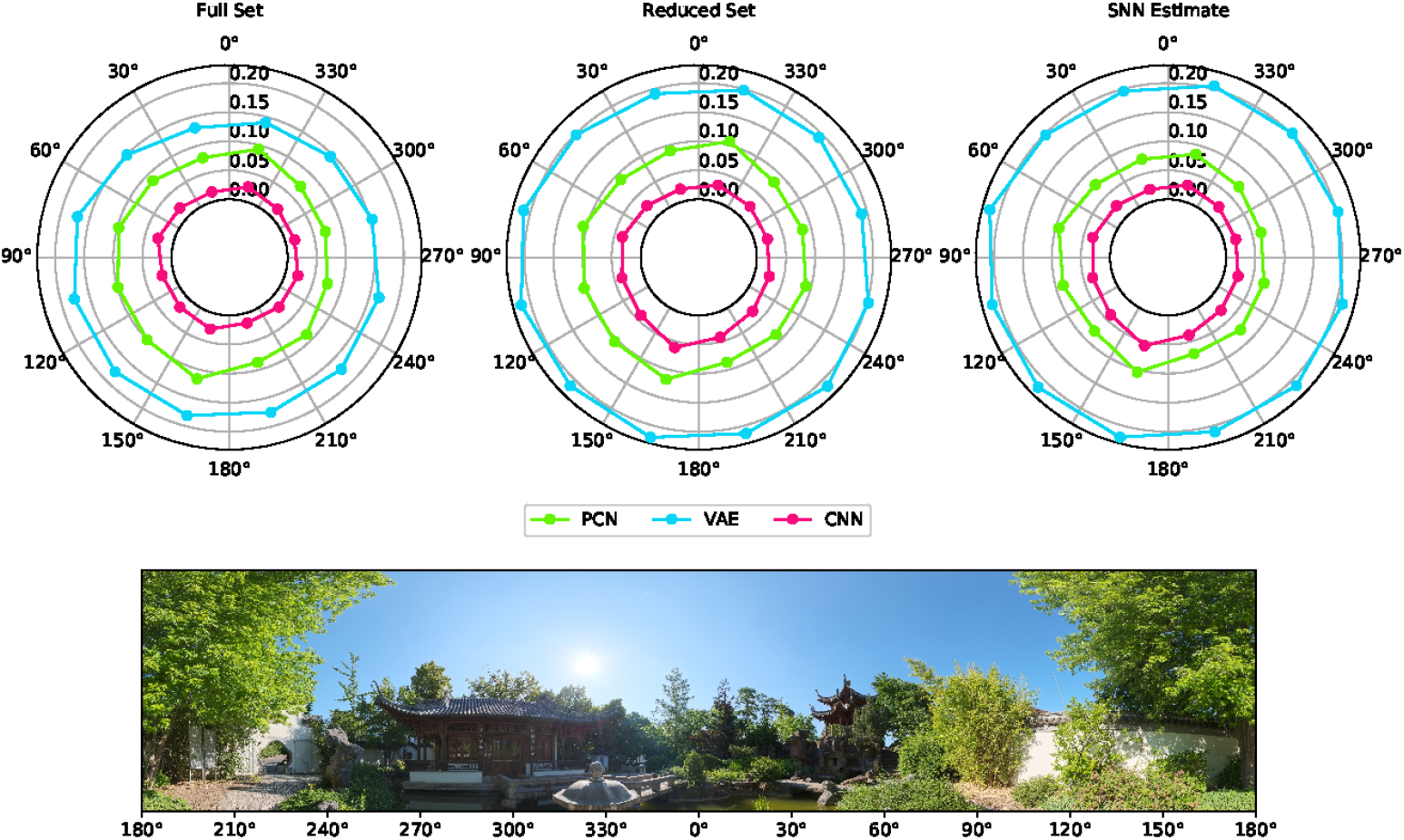
Reconstruction quality for different viewpoints of the environment taken from the rotating test dataset. The reconstruction quality for each 30 degree arc of view is represented as a point with radial distance proportional to the Root Mean Squared Error between the ground truth Laplacian and each model reconstruction (PCN, VAE and CNN trained using each of the training sets). The panorama depicts the view point of the robot as it rotates on the spot with associated angular head direction labeled in register with the error polar plots above.

Figure 7 shows the overall reconstruction quality for all datasets and scenarios. For all three models, reconstruction quality was noticeably impacted by a reduction in dataset quality, but the absolute error remains small. Both the reduced dataset and the SNN estimates were comparable in their error values, demonstrating that the internally generated estimates of the SNN model are a suitable substitute for ground truth odometry as a teaching signal, provided the dataset (and therefore accumulated drift) is small. Further to this, it demonstrates the effectiveness of all three methods, and thus their representative paradigms, at performing this task with limited data.

**Figure 7:**
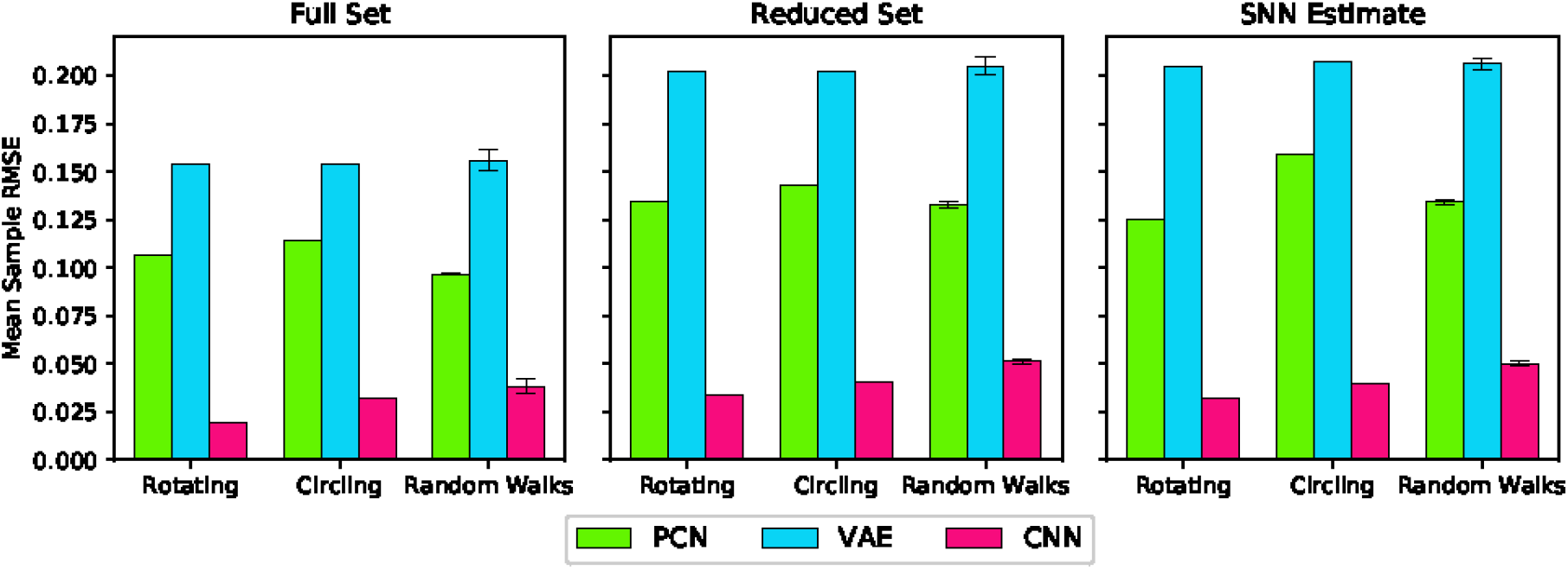
Reconstruction quality (Root Mean Squared Error) for each model during each behaviour trained using each training set. Random Walks columns represent the mean of the 5 Random Walk datasets with error bars indicating the standard error.

### Drift reduction using head direction predictions as allothetic input

The predictions generated by the PCN, VAE and CNN were converted into one-to-one current inputs to LMN cells to correct for drift using visual information. Figure 8 shows the ground truth head direction, ideothetic only estimate and the corrected estimate for 3 example datasets (rotation, a random walk, and circling), with the respective error over time. In each case the corrected head direction estimate (pink) is much closer to the ground truth (grey) than the estimate using ideothetic input only (blue) which drifts over time. Across all five random walk datasets, corrective input from the PCN, VAE and CNN all significantly reduced drift (one way ANOVA with Tukey HSD post-hoc testing: PCN p=0.001; VAE p¡0.001; CNN p¡0.001, Table 2). The smallest error after corrections was achieved using predictions made by the CNN, which had the best reconstruction quality. Even though VAE prediction is imprecise it still performs comparably to the other methods. This may be due to current inputs onto HD cells far from the active bump having less influence due to attractor dynamics. Only current inputs close to the bump location have strong influence over bump position. Although the drift was large for the circling dataset (RMSE = 9.75 radians), all three methods successfully corrected for this drift. This was the biggest reduction in error for all three model-free learning algorithms (difference in RMSE, PCN 9.59 radians, VAE 9.69 radians, CNN 9.71 radians).

**Table 2:**
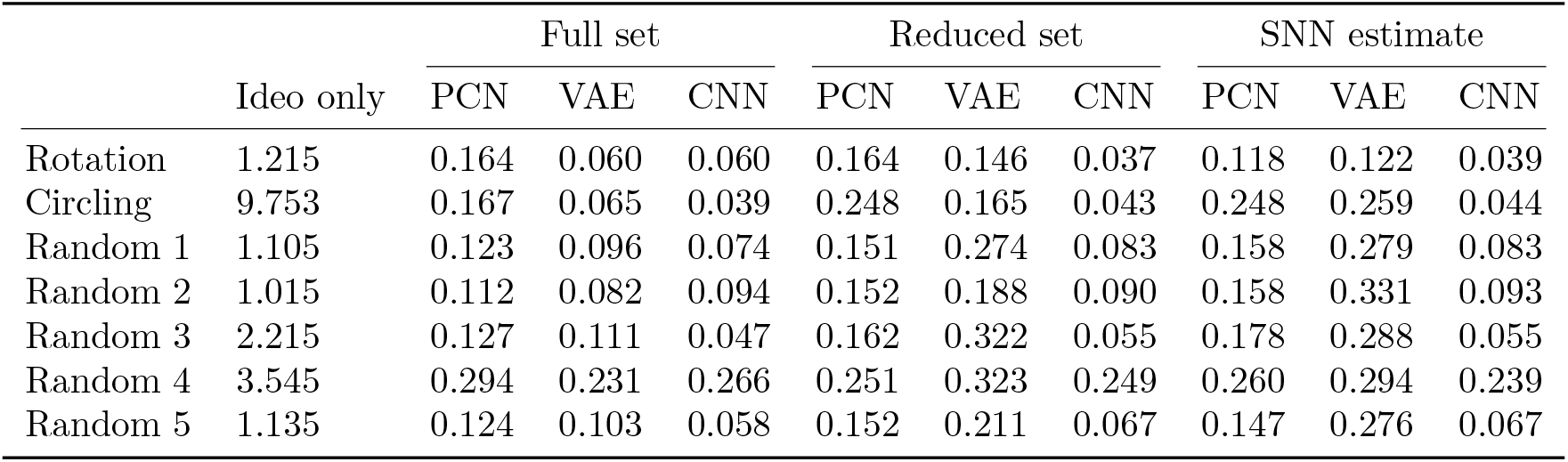
RMSE of the difference between the estimated head direction from the model and the ground truth using only ideothetic drive, and with corrections from the PCN, VAE or CNN trained on each of the three training sets.

**Figure 8:**
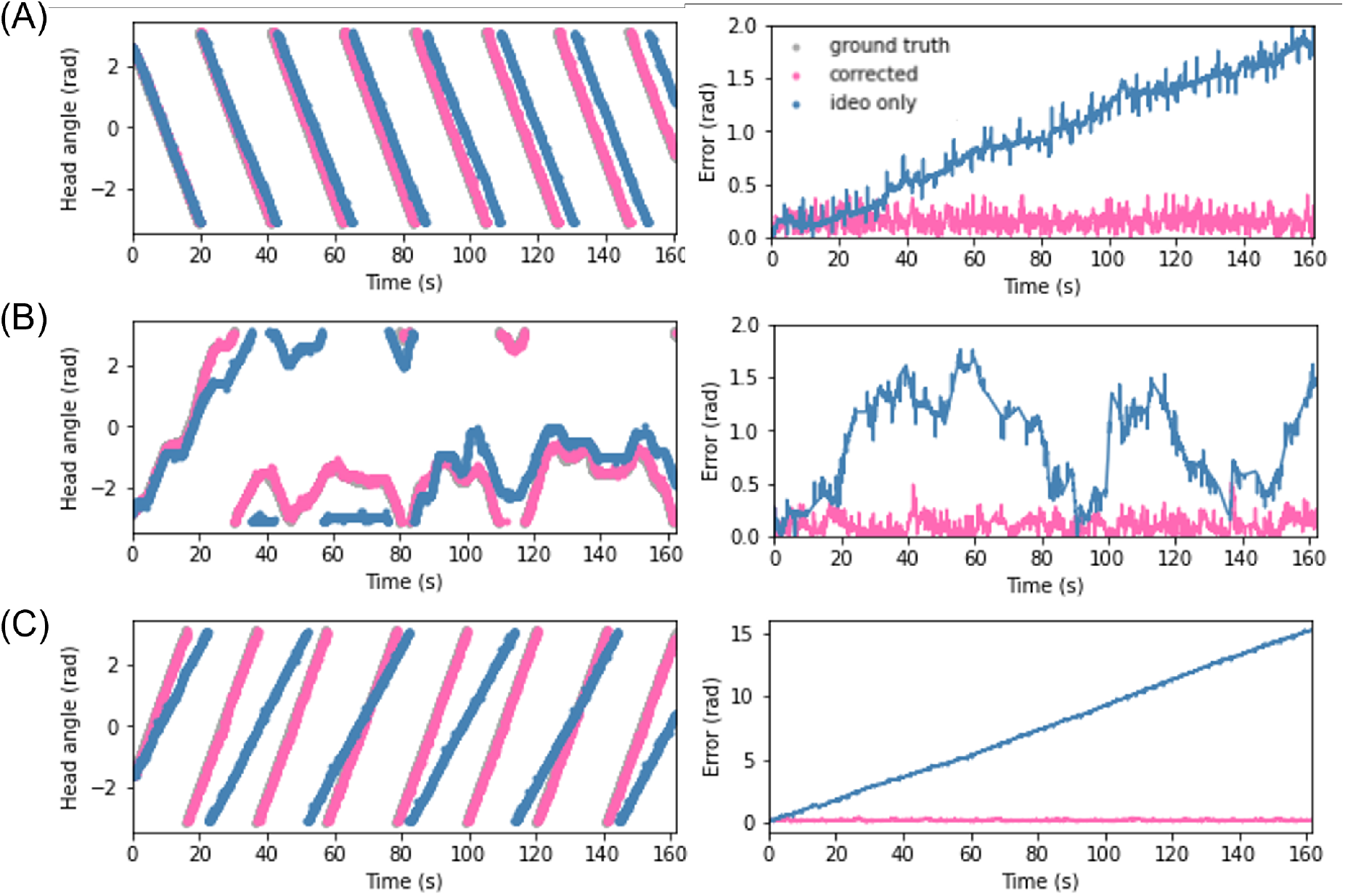
Plots showing estimated head angle from the SNN with ideothetic drive only (blue), the corrected estimated head angle from the SNN, and ground truth head angle (grey) which also receives allothetic input from the PCN (left). The difference between each estimate and the ground truth as also show (right). Examples are shown from the rotating **(A)**, random walk 1 **(B)** and circling **(C)** datasets. In all cases the allothetic correction results in minimised drift and the corrected estimate and ground truth are almost indistiguishable. As the PCN, VAE and CNN produce similar reductions in drift, only the PCN plots are shown.

### Drift reduction using a reduced training set

Ground truth head direction is not available in biology to form associations between visual scenes and heading. As drift in the head direction estimate is minimal (RMSE) during the first rotation when only idiothetic information is available, this estimated head direction could be used for training the model-free learning algorithms. This would be a much smaller training set, so we first used a single rotation of the ground truth to test the PCN, VAE and CNNs ability to predict head direction from a smaller training set. The PCN, VAE and CNN, were trained using a single rotation of ground truth head directions, and the same method was used to convert the predictions into current input to the head direction cells. In all cases the RMSE between ground truth and the estimate head direction was reduced. Across all five random walk datasets, corrective input from the PCN, VAE and CNN all significantly reduced drift (one way ANOVA with Tukey HSD post-hoc testing: PCN p=0.001; VAE p=0.002; CNN p¡0.001, Table 2). Once again the largest error reductions were achieved using CNN predictions. The VAE corrections were the least helpful, reflecting the lower reconstruction quality achieved training on the small dataset.

### Drift reduction using SNN estimate as training set

As the reduction in drift was comparable when the full 3 minute ground truth and a single revolution were used as training sets, we trained each of the model-free learning algorithms on a single revolution of the estimated head direction produced by the spiking model.

Similar to the drift reduction seen for the previous two training sets. Drift was reduced by all three models from all of the datasets. Across all five random walk datasets, corrective input from the PCN, VAE and CNN all significantly reduced drift (one way ANOVA with Tukey HSD post-hoc testing: PCN p=0.001; VAE p=0.002; CNN p¡0.001, Table 2). CNN produces the best error reduction, ahead of the PCN and then VAE, reflecting the reconstruction quality of their predictions. With each decrease in training set quality from full ground truth, to single revolution ground truth to single revolution estimated head direction the average error across the random walks increased for both the PCN and VAE, remaining stable only for the CNN. Figure 9 shows a summary of drift reduction by all three model-free learning algorithms trained on the full, reduced and SNN estimate training sets. Compared to head direction estimates which rely only on ideothetic input, or randomly generated predictions, all methods and training sets produced a significant reduction in drift.

**Figure 9:**
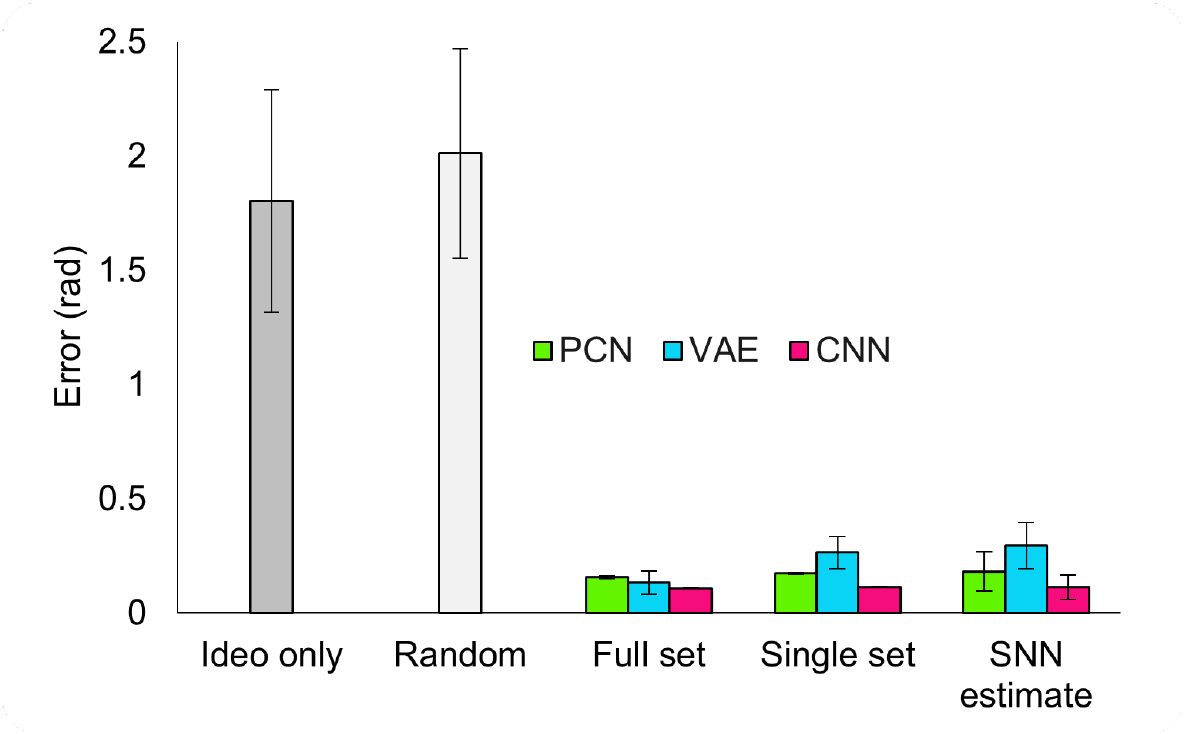
Summary of drift reduction for each of the model-free learning algorithms and training sets across all 5 random walk datasets. Compared to idiothetic input only and random predictions, drift is significantly reduced for all three methods trained on full, single and SNN estimate training sets. Bar plot shows average error (radians) *±* standard error.

### Cue rotation

Head direction cells in rodent have been shown to follow environmental cues over their ideothetic estimate of heading, even when the cues are rotated within the environment ([Taube and Burton, 1995, Yoder et al., 2015]). To replicate a cue rotation experiment using the WhiskEye rotating behaviour, we provided raw allothetic predictions from each of the model-free learning algorithms for the first minute, then rotated *π*/2 radians either clockwise or anticlockwise for 30 seconds before returning to raw allothetic predictions. Figure 10 shows the head direction estimate against the ground truth, with the error for clockwise (Figure 10A) and anti-clockwise (Figure 10B) rotations of the allothetic input from the PCN, VAE and CNN trained on the full ground truth. The red line shows an error of *π*/2 radians, which is the rotation of the cue.

**Figure 10:**
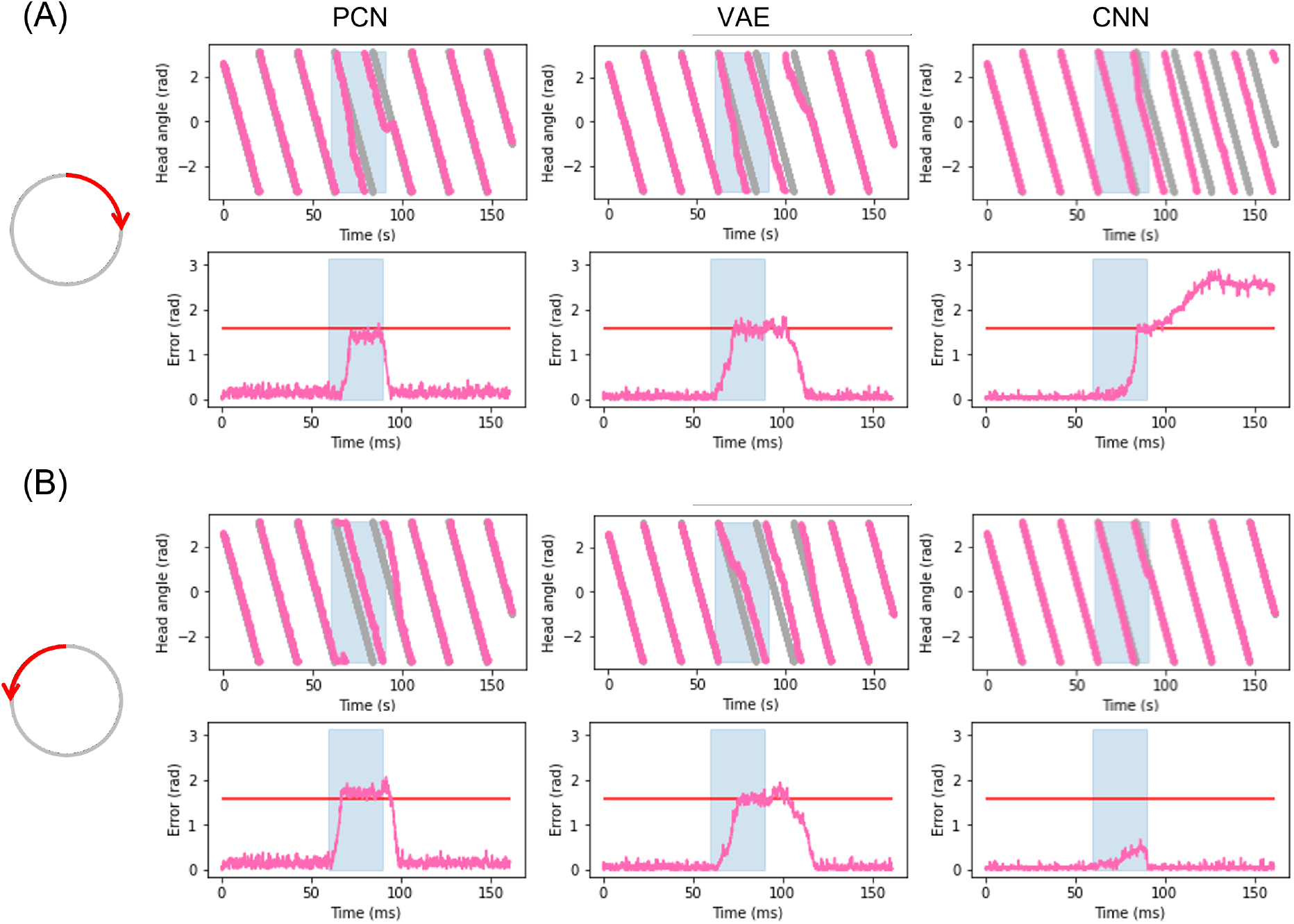
Plots showing corrected estimated head angle (pink) compared the ground truth (grey) during the artificial cue rotation experiments. Blue block indicates period of cue rotation either clockwise **(A)** or anticlockwise **(B)**, and the expected rotation (*π*/2 radians) is indicated with a red line on the error plot. In both clockwise and anticlockwise rotations corrections by the PCN and the VAE move the bump to the rotated position after a delay. The CNN fails to pull the bump contrary to the direction of bump movement and cannot maintain control over bump position in the clockwise rotation.

For both clockwise and anti-clockwise rotations, the PCN and VAE input strongly control the bump position. After a short delay, the bump position moves the full *π*/2 radians, error between ground truth and the estimate reaching the red line. When the rotation is removed the bump continues to follow the allothetic input after a delay. Some drift may be required before the allothetic input can gain control over the bump position, resulting in a delay. In the case of the CNN, only when the ideothetic drive and the rotation were in the same direction (Figure 10A) could the allothetic input control the bump position strongly enough to complete the full rotation. The allothetic input cannot pull the bump back round in the opposite direction to the ideothetic drive when the rotation is removed, resulting in drift as if no allothetic input is available. Because the allothetic and ideothetic input are provided simultaneously, the bump is more likely to move when both of these pull the bump in the same direction around the ring rather than compete with each other.

The CNN has consistently the highest reconstruction quality of all three methods (Figure 5), producing predictions with a sharp Laplacian peak. This prediction shape results in current input to a small number of cells at a precise position, and produces the most accurate head direction estimate. This is likely because the amount of drift between each allothetic correction is small, and the bump does not need to be moved far, noisier predictions from the VAE and PCN result in current injection to more cells, making it less accurate from drift correction but more able to move the bump large distances like in this cue conflict case. These data suggest a refined Laplacian peak is not the most effective prediction shape for strong allothetic control over the head direction signal. In all cases the current magnitude used was high enough to correct for drift without impairing ideothetic control. By varying the amount of current supplied, allothetic input could have stronger or weaker control over the bump position regardless of prediction shape.

## Discussion

With these experiments we have shown that, like head direction cells recorded in rodents, a spiking continuous attractor model of head direction cells driven purely by self motion (ideothetic) information is subject to drift. Taking inspiration from a number of previous studies ([Boucheny et al., 2005, Song and Wang, 2005, Shipston-Sharman et al., 2016]), we exploit reciprocal excitatory and inhibitory connections between the LMN and DTN to produce attractor dynamics which maintain a bump of activity at the estimated head angle.

Drift is thought to be caused by imprecise self motion cues but may also be due to inaccuracies in the model of angular head velocity (AHV). Variability within environments or body, such as injury (or in robots; inaccuracy in the odometry data due to wheel slip) make maintaining a precise model of AHV at all times unlikely. A prominent limitation of the experimental apparatus is that the odometry from the robot being collected from a simulated embodiment is not subject to inaccuracies. However, the stochastic nature of a spiking model limits the resolution and range of angular velocities which can be accurately represented by a single neuron, this can be seen clearly in the large drift accrued during the circling dataset where head angle changes very slowly. Using a population code rather than single cells may allow for a finer resolution of AHVs which can be represented in spikes, and contribute to reducing drift.

In rodents, drift in the preferred firing directions of head direction cells is seen mainly in the dark or when brain regions providing allothetic input are lesioned (primarily visual; [Yoder et al., 2015]) indicating these data are essential for stabilising the head direction signal. Using predictions from three different model-free learning algorithms, we directly influenced the bump position, minimising drift. In some previous models of drift correction, allothetic information contributes to calibrating the model of AHV, rather than using allothetic input to directly change the bump position. Kreiser et al. (2020) refine the AHV model by detecting error between the estimated head angle and learnt positions of landmarks, and altering firing properties of AHV cells. Stratton et al (2011) suggest a role for specific behaviour patterns for learning new landmarks and calibrating the AHV model. We show that predictions made after training on the estimated head angle from the SNN during a single revolution (specific behaviour), can be used to successfully correct for drift.

Entrainment of the head direction signal to visual information has been seen in cue rotation studies, where external environmental cues are rotated in the environment and a corresponding rotation is observed in the preferred firing direction of the head direction cells. These large changes in bump location are better solved by influencing the bump position directly, rather than updating the AHV model. By rotating the allothetic predictions, we have replicated shifts in the bump position to match the rotation of the environment. As AHV cell firing also shows some refinement when visual information is available ([Keshavarzi et al., 2021]), going forward a combination of optimising the AHV model and direct bump movement could be used.

A Laplacian-shaped input centred on the current HD, was used to train the three model-free learning algorithms. The CNN reproduced this shape in its prediction whereas the VAE and PCN produced broader more Gaussian shaped predictions. The CNN consistently produced the most precise head direction predictions even for the small SNN estimate training set. This suggests a trivial learning problem for the CNN, likely because the range of possible distal views observed by the robot is small and bounded; the same frames used for training the CNN are likely to be reobserved as the robot rotates. However even with less precise predictions the PCN and VAE can reduce drift significantly, likely due to the attractor dynamics dampening current inputs far from the bump location. The artificial cue conflict experiments revealed a precise Laplacian distribution not to be suitable as a corrective signal due to the limited number of cells current is injected into, and therefore the limited power of this input to influence the bump location. The broader predictions made by the PCN and the VAE were able to better control the bump position. Refining the shape and strength of the prediction would likely change the allothetic control over the bump. Two populations of head direction cell have been identified in rodents, those more controlled by allothetic input and others more strongly controlled by ideothetic input ([Dudchenko et al., 2019]). We can see how by varying the strength or shape of allothetic input to the network, these two cell types may emerge.

In previous work, correcting drift with the aid of visual information has either assumed visual processing upstream and provided correction based on the ground truth ([Song and Wang, 2005]), or learnt the orientation of arbitrary features, such as LEDs or coloured panels ([Kreiser et al., 2020, Yan et al., 2021]). Here we show that corrective signals can be generated by learning associations between natural visual scenes and a self generated representation of heading, without identifying specific environmental landmarks. However we recognise, including advanced visual processing and feature extraction may be useful for online learning mechanisms to determine the reliability of visual input. This type of corrective allothetic signal is presumed to come from the postsubiculum. Lesions of this region lead to more drift than seen for just control animals in the dark ([Yoder et al., 2015]). This suggests this region may be contributing more than just visual correction, but also other sensory modalities. In this paper we have focused on the calibration of head direction estimate by visual inputs, however an intriguing direction for future work would be the inclusion of other allothetic information such as tactile or olfactory. In visually ambiguous environments, conflicting visual cues may cause the HD estimate to become less accurate. Olfaction has great potential for detecting loop closures, as rodents leave scent trails as the explore environments ([Peden and Timberlake, 1990]), which can tell them if and how long ago they visited a position. A recent study in mice has shown that blind animals can use olfactory information to correct for drift in the head direction estimate ([Asumbisa et al., 2022]).

All of these methods currently require batch learning of head direction image pairs, however, as rodents continue to move within environments they must continually learn and refine associations between head angle and visual scenes. Learning to place less weight on unreliable cues such as the position of the sun, which may initially appear as a useful landmark but becomes unstable with time ([Knight et al., 2014]), is key to reliable correction of head angle in dynamic natural environments. The next step is to adapt these model-free learning algorithms to learn continuously and adapt its predictions as the robot explores its environment.

The three trained models, despite their differences in reconstruction quality and overall error, are all good candidates for generating allothetic corrections for the SNN. Although some scenarios such as cue conflicts show weakness of overly-precise estimates as by the CNN, this is not a fault of the model itself; the robustness of the PCN and VAE predictions to cue conflicts shows that RMSE from a Laplacian ground truth signal is not ideal for the task at hand, and that better performance could be gained by training to a broader distribution (such as a Gaussian).

Where differences do lie is in their applicability to more complex experimental setups. The environment the data is gathered from is simple in structure despite the complexity of the visual scene; there are no proximal cues to obscure the environment and sensory input is limited to vision. Previous studies have shown non visual and multimodal examples of CNNs ([Dauphin et al., 2017, Ma et al., 2015]) and VAE architectures ([Suzuki et al., 2017]) can perform well. Both, however, have issues with scaling: the multimodal CNN requiring many stacked networks working together, and multimodal VAEs requiring many intermediate uni-modal latent spaces to perform the task successfully. It is an open question as to how well PCNs will scale into more than 2 modalities and whether they will run into similar scaling issues as the VAEs. However, its method of operation and learning rule are bio-plausible, with local learning making it the best candidate for implementation as a spiking model. Furthermore, prior work has already shown that a PCN can use tactile information to inform localisation ([Pearson et al., 2021]).

This work has raised many important questions. How robust are these model-free learning approaches to a changing world, particularly with multiple environments, visually and potentially tactually distinct from each other? How can these be trained in a sequential manner, as an animal would experience them, whilst avoiding the catastrophic forgetting of earlier environments? Can the bio-plausibility of this system be increased by making the learning fully online, and is the SNN estimate of head direction reliable for long enough to train one of these algorithms to produce useful corrections? The experimental apparatus developed and used in this study are well placed to address these questions.

## Conclusion

Ideothetic control of the head direction system is especially important in new, ambiguous or dark environments, and allothetic control increases the accuracy of the head direction estimate and may help refine ideothetic control or make large corrections after a period of drift. We have shown that natural visual scenes, without identifying specific landmarks, can be used to predict the current head angle by training three model-free learning algorithms on a limited and imprecise training set of estimated head angles produced by a spiking continuous attractor model of the head direction cell system driven by ideothetic inputs from robot odometry. Predictions from all three methods were equally valuable in minimising drift.

## Acknowledgments

This research has received funding from the European Union’s Horizon 2020 Framework Programme for Research and Innovation under the Specific Grant Agreement No. 945539 (Human Brain Project SGA3).

Chinese Garden in Stuttgart panorama taken and released under CC0 license by Andreas Mitchok.

## References

K. Asumbisa, A. Peyrache, and S. Trenholm. Flexible cue anchoring strategies enable stable head direction coding in both sighted and blind animals. bioRxiv, page 2022.01.12.476111, 1 2022. doi: 10.1101/2022.01.12.476111.

J. P. Bassett and J. S. Taube. Neural correlates for angular head velocity in the rat dorsal tegmental nucleus. The Journal of neuroscience : the official journal of the Society for Neuroscience, 21:5740–51, 8 2001.

J. P. Bassett, T. J. Wills, and F. Cacucci. Self-organized attractor dynamics in the developing head direction circuit. Current Biology, 28:609–615.e3, 2 2018. doi: 10.1016/j.cub.2018.01.010.

Y. Bengio and Y. Lecun. Convolutional networks for images, speech, and time-series. 11 1997.

A. Bicanski and N. Burgess. Environmental anchoring of head direction in a computational model of retrosplenial cortex. Journal of Neuroscience, 36:11601–11618, 11 2016. doi: 10.1523/JNEUROSCI.0516-16.2016.

H. T. Blair, J. Cho, and P. E. Sharp. The anterior thalamic head-direction signal is abolished by bilateral but not unilateral lesions of the lateral mammillary nucleus. The Journal of neuroscience : the official journal of the Society for Neuroscience, 19:6673–83, 8 1999.

C. Boucheny, N. Brunel, and A. Arleo. A continuous attractor network model without recurrent excitation: maintenance and integration in the head direction cell system. Journal of computational neuroscience, 18:205–227, 3 2005. doi: 10.1007/S10827-005-6559-Y.

L. Breiman, J. H. Friedman, R. A. Olshen, and C. J. Stone. In Classification and Regression Trees, 1983.

J. Cho and P. E. Sharp. Head direction, place, and movement correlates for cells in the rat retrosplenial cortex. Behavioral Neuroscience, 115:3–25, 2001. doi: 10.1037/0735-7044.115.1.3.

F. Chollet et al. Keras. https://keras.io, 2015.

Y. N. Dauphin, A. Fan, M. Auli, and D. Grangier. Language modeling with gated convolutional networks. In Proceedings of the 34th International Conference on Machine Learning - Volume 70, ICML’17, page 933–941. JMLR.org, 2017.

J. J. DiCarlo, R. Haefner, L. Isik, T. Konkle, N. Kriegeskorte, B. Peters, N. Rust, K. Stachenfeld, J. B. Tenenbaum, D. Tsao, and I. Yildirim. How does the brain combine generative models and direct discriminative computations in high-level vision? In Generative Adversarial Collaborations, 2021.

S. Dora, C. Pennartz, and S. Bohte. A deep predictive coding network for inferring hierarchical causes underlying sensory inputs. In K. V., M. Y., H. B., I. L., and M. I., editors, Artificial Neural Networks and Machine Learning – ICANN 2018., volume 11141. Springer, 2018. doi: 10.1007/978-3-030-01424-745.

P. A. Dudchenko, E. R. Wood, and A. Smith. A new perspective on the head direction cell system and spatial behavior. Neuroscience and Biobehavioral Reviews, 105:24–33, 10 2019. doi: 10.1016/j.neubiorev.2019.06.036.

E. Falotico, L. Vannucci, A. Ambrosano, U. Albanese, S. Ulbrich, J. C. V. Tieck, G. Hinkel, J. Kaiser, I. Peric, O. Denninger, N. Cauli, M. Kirtay, A. Roennau, G. Klinker, A. V. Arnim, L. Guyot, D. Peppicelli, P. Martínez-Cañada, E. Ros, P. Maier, S. Weber, M. Huber, D. Plecher, F. Röhrbein, S. Deser, A. Roitberg, P. van der Smagt, R. Dillman, P. Levi, C. Laschi, A. C. Knoll, and M.-O. Gewaltig. Connecting artificial brains to robots in a comprehensive simulation framework: The neurorobotics platform. Frontiers in Neurorobotics, 11, 1 2017. ISSN 1662-5218. doi: 10.3389/fnbot.2017.00002.

I. J. Goodfellow, J. Pouget-Abadie, M. Mirza, B. Xu, D. Warde-Farley, S. Ozair, A. Courville, and Y. Bengio. Generative adversarial networks, 2014.

J. P. Goodridge and J. S. Taube. Preferential use of the landmark navigational system by head direction cells in rats. Behavioral neuroscience, 109:49–61, 1995. doi: 10.1037/0735-7044.109.1.49.

J. P. Goodridge, P. A. Dudchenko, K. A. Worboys, E. J. Golob, and J. S. Taube. Cue control and head direction cells. Behavioral neuroscience, 112:749–761, 1998. doi: 10.1037//0735-7044.112.4.749.

R. M. Grieves and K. J. Jeffery. The representation of space in the brain. Behavioural processes, 135: 113–131, 2 2017. doi: 10.1016/J.BEPROC.2016.12.012.

T. Hafting, M. Fyhn, S. Molden, M.-B. Moser, and E. I. Moser. Microstructure of a spatial map in the entorhinal cortex. Nature, 436:801–806, 8 2005. doi: 10.1038/nature03721.

J. Hawkins, M. Lewis, M. Klukas, S. Purdy, and S. Ahmad. A framework for intelligence and cortical function based on grid cells in the neocortex. Frontiers in Neural Circuits, 12, 2019. doi: 10.3389/fn-cir.2018.00121.

K. He, X. Zhang, S. Ren, and J. Sun. Deep residual learning for image recognition. In 2016 IEEE Conference on Computer Vision and Pattern Recognition (CVPR), pages 770–778, 2016. doi: 10.1109/CVPR.2016.90.

S. Keshavarzi, E. F. Bracey, R. A. Faville, D. Campagner, A. L. Tyson, S. C. Lenzi, T. Branco, and T. W. Margrie. Multisensory coding of angular head velocity in the retrosplenial cortex. Neuron, 11 2021. doi: 10.1016/J.NEURON.2021.10.031.

D. P. Kingma and M. Welling. Auto-encoding variational bayes. In ICLR 2014 Workshop, 2013.

R. Knight, C. E. Piette, H. Page, D. Walters, E. Marozzi, M. Nardini, S. Stringer, and K. J. Jeffery. Weighted cue integration in the rodent head direction system. Philosophical Transactions of the Royal Society B: Biological Sciences, 369, 2 2014. doi: 10.1098/rstb.2012.0512.

T. C. Knowles, R. Stentiford, and M. J. Pearson. Whiskeye: A biomimetic model of multisensory spatial memory based on sensory reconstruction. In C. Fox, J. Gao, A. Ghalamzan, E. M. Saaj, M. Hanheide, and S. Parsons, editors, Lecture Notes in Artificial Intelligence 13054, pages 408–418. Springer, 9 2021. doi: 10.1007/978-3-030-89177-043.

R. Kreiser, G. Waibel, N. Armengol, A. Renner, and Y. Sandamirskaya. Error estimation and correction in a spiking neural network for map formation in neuromorphic hardware. 2020 IEEE International Conference on Robotics and Automation (ICRA), pages 6134–6140, 5 2020. doi: 10.1109/ICRA40945.2020.9197498.

E. Kropff, J. E. Carmichael, M. B. Moser, and E. I. Moser. Speed cells in the medial entorhinal cortex. Nature, 523:419–424, 7 2015. doi: 10.1038/nature14622.

C. Lever, S. Burton, A. Jeewajee, J. O’Keefe, and N. Burgess. Boundary vector cells in the subiculum of the hippocampal formation. The Journal of neuroscience : the official journal of the Society for Neuroscience, 29:9771–9777, 8 2009. doi: 10.1523/JNEUROSCI.1319-09.2009.

L. Ma, Z. Lu, L. Shang, and H. Li. Multimodal convolutional neural networks for matching image and sentence. 2015 IEEE International Conference on Computer Vision (ICCV), pages 2623–2631, 2015.

B. L. McNaughton, F. P. Battaglia, O. Jensen, E. I. Moser, and M. B. Moser. Path integration and the neural basis of the ’cognitive map’. Nature Reviews Neuroscience 2006 7:8, 7:663–678, 8 2006. doi: 10.1038/nrn1932.

J. O’Keefe. Place units in the hippocampus of the freely moving rat. Experimental Neurology, 51:78–109, 1976. doi: 10.1016/0014-4886(76)90055-8.

H. J. Page and K. J. Jeffery. Landmark-based updating of the head direction system by retrosplenial cortex: A computational model. Frontiers in Cellular Neuroscience, 12:191, 7 2018. doi: 10.3389/FN-CEL.2018.00191/BIBTEX.

M. Pearson, S. Dora, O. Struckmeier, T. Knowles, B. Mitchinson, K. Tiwari, V. Kyrki, S. Bohte, and C. Pennartz. Multimodal representation learning for place recognition using deep hebbian predictive coding. Frontiers in Robotics and AI, 8:403, 2021. doi: 10.3389/frobt.2021.732023.

B. F. Peden and W. Timberlake. Environmental influences on flank marking and urine marking by female and male rats (rattus norvegicus). Journal of comparative psychology (Washington, D.C. : 1983), 104: 122–130, 1990. doi: 10.1037/0735-7036.104.2.122.

D. A. Reynolds. Gaussian mixture models. In Encyclopedia of Biometrics, 2009.

P. E. Sharp, A. Tinkelman, and J. Cho. Angular velocity and head direction signals recorded from the dorsal tegmental nucleus of gudden in the rat: Implications for path integration in the head direction cell circuit. Behavioral Neuroscience, 115:571–588, 2001. doi: 10.1037/0735-7044.115.3.571.

O. Shipston-Sharman, L. Solanka, and M. F. Nolan. Continuous attractor network models of grid cell firing based on excitatory–inhibitory interactions. Journal of Physiology, 594:6547–6557, 11 2016. doi: 10.1113/JP270630.

P. Song and X. J. Wang. Angular path integration by moving “hill of activity”: A spiking neuron model without recurrent excitation of the head-direction system. Journal of Neuroscience, 25:1002–1014, 1 2005. doi: 10.1523/JNEUROSCI.4172-04.2005.

R. W. Stackman and J. S. Taube. Firing properties of rat lateral mammillary single units: head direction, head pitch, and angular head velocity. The Journal of neuroscience : the official journal of the Society for Neuroscience, 18:9020–37, 11 1998.

R. W. Stackman, E. J. Golob, J. P. Bassett, and J. S. Taube. Passive transport disrupts directional path integration by rat head direction cells. Journal of neurophysiology, 90:2862–2874, 11 2003. doi: 10.1152/JN.00346.2003.

P. Stratton, M. Milford, G. Wyeth, and J. Wiles. Using strategic movement to calibrate a neural compass: A spiking network for tracking head direction in rats and robots. PLoS ONE, 6:25687, 2011. doi: 10.1371/journal.pone.0025687.

M. Suzuki, K. Nakayama, and Y. Matsuo. Joint multimodal learning with deep generative models. In ICLR 2017 Workshop, 2017.

J. S. Taube. Head direction cells recorded in the anterior thalamic nuclei of freely moving rats. The Journal of neuroscience : the official journal of the Society for Neuroscience, 15:70–86, 1 1995.

J. S. Taube and H. L. Burton. Head direction cell activity monitored in a novel environment and during a cue conflict situation. Journal of neurophysiology, 74:1953–1971, 1995. doi: 10.1152/JN.1995.74.5.1953.

J. S. Taube, R. U. Muller, and J. B. Ranck. Head-direction cells recorded from the postsubiculum in freely moving rats .1. description and quantitative-analysis. Journal of Neuroscience, 10:420–435, 1990.

S. D. Vann, J. P. Aggleton, and E. A. Maguire. What does the retrosplenial cortex do? Nature Reviews Neuroscience, 10:792–802, 11 2009. doi: 10.1038/nrn2733.

Y. Yan, N. Burgess, and A. Bicanski. A model of head direction and landmark coding in complex environments. PLoS computational biology, 17, 9 2021. doi: 10.1371/JOURNAL.PCBI.1009434.

R. M. Yoder and J. S. Taube. Head direction cell activity in mice: robust directional signal depends on intact otolith organs. The Journal of neuroscience : the official journal of the Society for Neuroscience, 29:1061–1076, 1 2009. doi: 10.1523/JNEUROSCI.1679-08.2009.

R. M. Yoder and J. S. Taube. The vestibular contribution to the head direction signal and navigation. Frontiers in Integrative Neuroscience, 8:32, 4 2014. doi: 10.3389/fnint.2014.00032.

R. M. Yoder, B. J. Clark, and J. S. Taube. Origins of landmark encoding in the brain. Trends in neurosciences, 34:561–571, 2011. doi: 10.1016/J.TINS.2011.08.004.

R. M. Yoder, J. R. Peck, and J. S. Taube. Visual landmark information gains control of the head direction signal at the lateral mammillary nuclei. Journal of Neuroscience, 35:1354–1367, 2015. doi: 10.1523/JNEUROSCI.1418-14.2015.

L. Yu, M. Nassar, S. A. Park, S. Sweigart, and E. Boorman. Do grid codes afford generalization and flexible decision-making? In Generative Adversarial Collaborations, 2021.

K. Zhang. Representation of spatial orientation by the intrinsic dynamics of the head-direction cell ensemble: a theory. Journal of Neuroscience, 16:2112–2126, 3 1996. doi: 10.1523/JNEUROSCI.16-06-02112.1996.

